# Long-term fish assemblages of the Ohio River: altered trophic and life history strategies with hydrologic alterations and landuse modifications

**DOI:** 10.1101/529974

**Authors:** Mark Pyron, Meryl C. Mims, Mario M. Minder, Robert C. Shields, Nicole Chodkowski, Caleb C. Artz

## Abstract

Long-term monitoring of species assemblages provides a unique opportunity to test hypotheses regarding environmentally-induced directional trajectories of freshwater species assemblages. We used 57 years of lockchamber fish rotenone and boat electrofishing survey data (1957-2014) collected by the Ohio River Valley Water Sanitation Commission (ORSANCO) to test for directional trajectories in taxonomy, trophic classifications, and life history strategies of freshwater fish assemblages in the Ohio River Basin. We found significant changes in taxonomic and trophic composition of freshwater fishes in the Ohio River Basin. Annual species richness varied from 31 to 90 species and generally increased with year. Temporal trajectories were present for taxonomic and trophic assemblages. Assemblage structure based on taxonomy was correlated with land use change (decrease in agriculture and increase in forest). Taxonomic assemblage structure was also correlated with altered hydrology variables of increased minimum discharge, decreased fall rate, and increased rise rate. Trophic composition of fish catch correlated with land use change (decrease in agriculture and increase in forest) and altered hydrology. Altered hydrology of increased minimum discharge, increased fall discharge, decreased base flows, and increased number of high pulse events was correlated with increased counts of herbivore-detritivores and decreased counts of piscivores and planktivores. We did not find directional changes in life history composition. We hypothesized a shift occurred from benthic to phytoplankton production throughout the basin that may have decreased secondary production of benthic invertebrates. This may also be responsible for lower trophic position of invertivore and piscivore fishes observed in other studies.

## Introduction

Anthropogenic threats to freshwater ecosystems are numerous and globally widespread. Rivers are altered by multiple factors including watershed land use, hydrologic alterations, municipal and industrial effluent, and water withdrawals [1]. Conservation of water resources and increased water demands requires understanding historical and current effects of water and land use to help inform potential solutions via restoration or intervention [2]; key to this is the scale at which human activities affect biodiversity, and the patterns detectable across scales. Long-term monitoring data of biotic and abiotic factors, such as fish assemblages, provide an opportunity to test hypotheses regarding how environmental modifications affect the taxonomic, trophic, and life history compositions of aquatic organisms [3,4].

Land use changes from natural ecosystems to those dominated by intense agriculture, deforestation, or urbanization can dramatically alter fish biodiversity [5–7]. Modification from forest, grasslands, or wetlands to tillable agricultural land on a global scale provides numerous benefits to humans. However, these activities are often unsustainable [8]. When damaging agricultural practices are abandoned for best management or conservation goals, nutrient loading and pesticide contamination continue to affect stream ecosystems for years or decades, so-called the “ghost of land use past” [9].

The legacy of agriculture and land use is manifested in the Ohio River Basin, a watershed historically dominated by agriculture, but converted from agriculture to forest during the 1960s-80s [10]. The Ohio River Basin was drastically modified via logging and draining of wetlands following European colonization [11]. Urban and agricultural land use in the Ohio River basin were 13% and 74%, respectively, in 1930, exceeding critical threshold effects on water quality and aquatic health [10]. In addition, the Ohio River hosts invasive species including Common Carp (*Cyprinus carpio*), Silver Carp (*Hypophthalmichthys molitrix*), Bighead Carp (*Hypophthalmichthys nobilis*), and zebra mussels [12].

Despite these challenges, Thomas et al. [11] identified improvements in fish assemblage metrics, including species richness and an index of well-being, in the most recent 50 years that were correlated with higher water quality after implementation of the US Clean Water Act. However, the degree to which taxonomy, trophic classification, and life history strategies of fish assemblages throughout the basin have changed, and whether these changes are correlated with changes in land use and hydrology, remains unknown. Free of anthropogenic or other major disturbances, fish assemblages are expected to vary stochastically around an equilibrium structure, returning to average state with time over the long term [13,14]. Conversely, assemblages monitored over the long-term often exhibit directional trajectories in response to anthropogenic disturbance [15–17] or may transition to alternative community states [18].

Food webs depict trophic interactions between consumers and resources and allow analyses of community structure, stability, and ecosystem processes [19]. However, complex drivers of ecosystem variation and food web modification are difficult to identify [20]. For example, flow regimes influence freshwater food webs in complex and frequently contradicting manners. Hydrologic alterations due to construction of dams and levees in the Upper Mississippi River caused increased hydrogeomorphic complexity [21]. Increased complexity resulted in more diverse food sources for fish production, and increased variation in food sources for invertivore and piscivore fishes [22]. However, in the Ohio River, similar correlations between hydrologic alterations and changing food webs were not observed. Bowes [23] found that mean trophic positions of fishes decreased after dam construction, but trophic position was not correlated with hydrologic variables. Delong & Thoms [20] identified major changes in carbon sources for the Ohio River, and increased variation in mean nitrogen stable isotope ratios of fishes, following flow modifications that occurred from 1950-55. Bowes [23] identified carbon sources for Ohio River fishes collected from 1931-1970 and found that autochthonous algae was the major contributor. Food webs can also be influenced by biotic factors such as aquatic invasive species [24] that replace or supplement native species and can shift energy sources. We were interested if hydrologic alterations in the Ohio River resulted in modifications of trophic structure in fish assemblages.

Fish assemblages are often controlled by hydrologic regimes [25–27], considered the “master variable” for ecological integrity of lotic ecosystems [28]. Hydrologic regimes are a significant contributor to habitat variation, the habitat template (or “templet”, *sensu* Southwood [29]) upon which life history strategies evolve in response to environmental filtering [30]. Winemiller & Rose [31] proposed three general endpoint life history strategies for fishes based on generation time, fecundity, and juvenile survivorship – referred to as the trivariate continuum model. Equilibrium strategists are characterized by late maturity, low fecundity, and high juvenile survivorship; they are predicted to occur in stable environments. Periodic strategists are characterized by late maturity, high fecundity, and low juvenile survivorship; periodic stream fishes are predicted to occur in environments with predictable seasonal hydrology. Opportunistic strategists are characterized by early maturity, low fecundity, low juvenile survivorship; they are predicted to occur in harsh environments with unstable hydrology [32]. Mims & Olden [27] validated hydrology-life history expectations for fishes across the United States, finding that the proportion of opportunistic strategists increased with flow variability, the proportion of periodic strategists increased with hydrology seasonality, and equilibrium strategists were weakly associated with hydrologic variability and predictability. Perkin et al. [4] tested for temporal variation in Sabine River, USA, fish assemblages as a result of dam construction and subsequent hydrologic alteration. Built upon the trilateral continuum model applied by Mims & Olden [27], Perkin et al. [4] characterized species into four life history strategy categories: equilibrium, periodic, and opportunistic, as well as intermediate for species that are not strongly associated with one of the three endpoint strategies. Perkin et al. [4] found that dam construction resulted in reduced hydrologic variability and predicted changes in life history trait composition of the fishes: opportunistic strategist richness decreased and intermediate strategist richness was constant. We predicted similar patterns for life history trait composition in the Ohio River.

In the present study, we predicted basin-scale, long-term directional shifts in the taxonomic, trophic, and life history characteristics of fish assemblages of the Ohio River Basin. We tested for temporal changes by taxonomic classification, trophic traits, and life history strategies for fish assemblages. Combining taxonomic analyses with trait-based analyses links ecological functions to environmental variation at assemblage-level scales [33,34]. Our predictions were that assemblage composition varied predictably and directionally with hydrologic alterations caused by dams and land use variation such that 1) the fish assemblage will vary temporally with hydrologic alterations and land use change, 2) benthic invertivores decrease and planktivores increase with time, corresponding with the prevalence of algae in Ohio River foodwebs [23], and 3) variation in catch by life history strategies are correlated with hydrologic alteration.

## Materials and methods

### Fish assemblage and taxonomic data

The Ohio River Valley Water Sanitation Commission (ORSANCO) was founded in 1948 to monitor water quality and curb pollution throughout the Ohio River Basin (ORB). ORSANCO personnel surveyed the Ohio River and its tributaries via biological and chemical assessments. Biological assessments include July to October lockchamber rotenone surveys from 1957-2005 (www.orsanco.org). These surveys allow for analysis in long-term assemblage shifts for fish communities in the ORB [11]. Rotenone collections (377 surveys, 59 locations) were in lockchambers at dams located near the Mississippi River confluence (river km 10) upstream to lock #53 (Fig 1, river km 1,549.8), and are described in detail by Thomas et al. [11]. Boat electrofishing surveys (1864 surveys, 1362 locations) at 0.5 km reaches were conducted from 1990 – 2014 at river locations that differed from rotenone lockchamber collections. Our preliminary analyses of overlapping years of rotenone and electrofishing collections resulted in different assemblage structure (unpubl. results), thus we analyzed rotenone and electrofishing collections separately. In order to explore temporal patterns of variation in fish assemblages at the basin-scale, we aggregated data into annual CPUE counts for each taxon. CPUE units are number of individuals per lock survey or number of individuals per 0.5 km river distance electrofished. Several taxonomic groups were identified only to genus or family in the ORSANCO database (details in Thomas et al. [11]) and were not included in analyses (Appendix).

**FIG 1.**
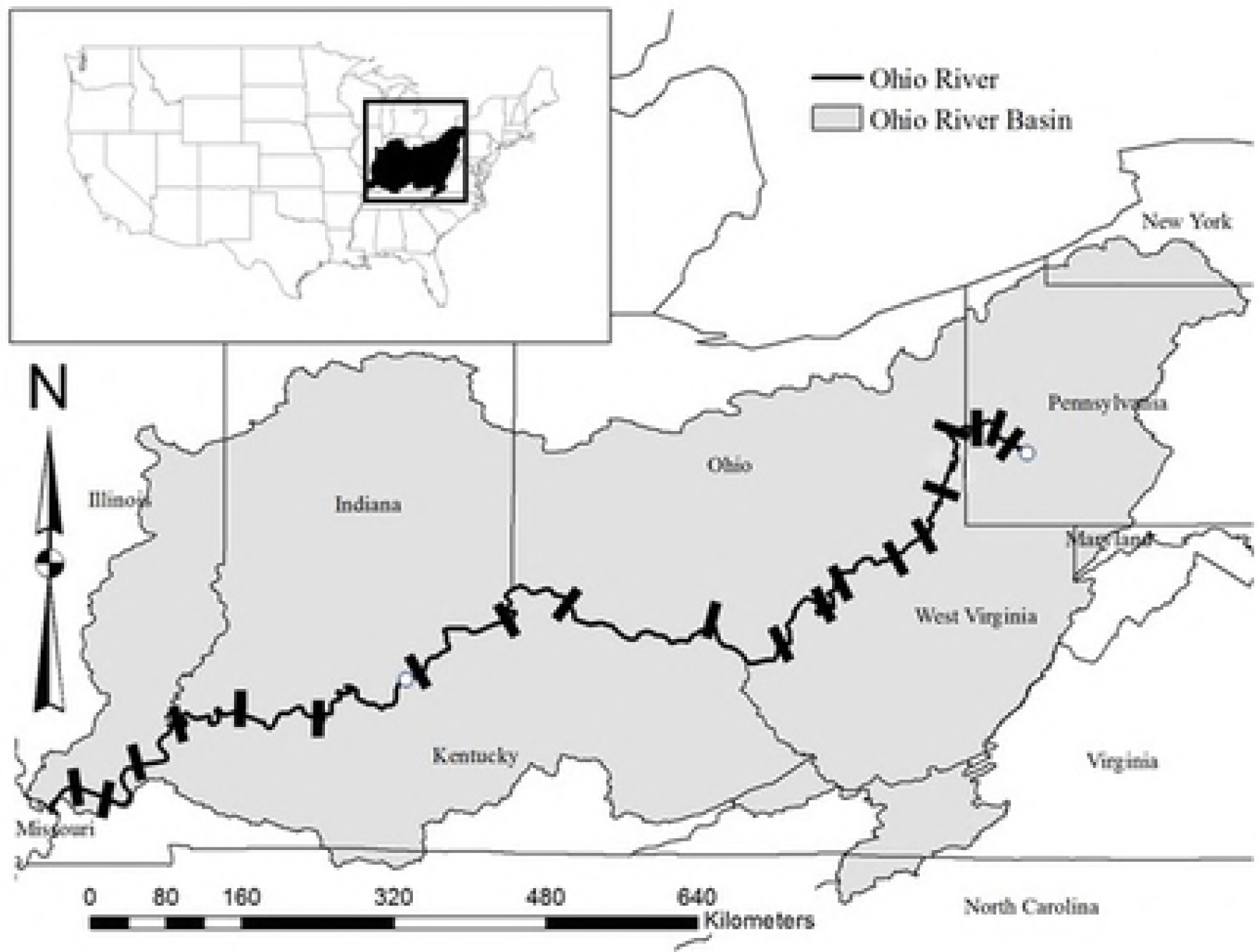
The Ohio River Basin showing locations of navigational dams (bars) and two USGS gaging stations (open circles). Locations for fish sampling are described by Thomas et al. [11]

### Environmental data

We used land use information for the Ohio River watershed from [35] that included 1957-2014 at a resolution of 0.5° × 0.5°. Land use was summarized by year and tested for correlation with year and for prediction of fish assemblages in ordinations (below). Impoundments were initially constructed over 100 years ago and have been frequently modified since [11]. We used daily discharge data from a downstream USGS site, at Louisville, OH (station number 03294500; referred to here as “Louisville”) and an upstream USGS site at Sewickley, PA (station number 03086000; referred to here as “Sewickley”) to analyze variation using the Indicators of Hydrologic Alteration (IHA) software [36]. We selected the default IHA parameters to identify altered variables, and for a single time period (same time period as fish data). Altered hydrology variables were defined by significant regressions with time (alpha = 0.05). We used a principal components analysis (PCA) to summarize dominant gradients of hydrologic variability as altered hydrology variables for each river location. We retained principal components (PC) with the largest eigenvalues from a scree plot, and included in constrained ordinations with CPUE counts of fishes by taxonomy, trophic traits, and life history strategies.

### Life history and trophic traits

Fishes were classified into the life history strategies defined by Winemiller & Rose [31] using the database originally published by Mims et al. [37] and recently modified to include updated traits data, including trophic information (Julian D. Olden, unpublished data). We calculated Euclidean distance in trivariate life history space from each species location to each of three life history strategy endpoints (opportunistic, periodic, equilibrium) and an intermediate strategy when species did not fit an endpoint (defined as strategy weights < 0.60), following Perkin et al. [4]. Euclidean distances were normalized between 0 and 1, and the inverse resulted in a “strategy weight” for each of the three life history gradients. We classified trophic traits (Julian D. Olden, unpublished data; [38]), which included adult feeding mode based on published diet analyses and we assigned each species to a trophic category (herbivore-detritivore, invertivore, invertivore-piscivore, omnivore, and piscivore).

### Analyses of basin-wide taxonomic, trophic, and life history composition through time

We used correlation analyses to examine species richness variation by year for CPUE counts by taxonomic, trophic, and life history categories. Additionally, we used a series of multivariate ordinations to explore taxonomic, trophic, and life history composition of fishes throughout the Ohio River Basin. Rare species (< 5 occurrences) were removed, and data were transformed by log (x + 1) or arcsine (percentage data). We used nonmetric multidimensional scaling (NMDS) with Bray-Curtis distance measure to explore directional shifts in the fish assemblage classified by taxonomy, trophic traits and life history strategy. We also used redundancy analysis (RDA), a constrained ordination, to test for relationships for hydrology and landuse variables with taxonomic, trophic, and life history composition of fish assemblages. RDA and NMDS analysis were performed in Canoco 5 [39], and NMDS ordination plots were created in R (R Development Core Team 2011). We used the forward selection of variables for a regression model option in Canoco, that uses a Monte Carlo permutation test for significance.

## Results

Long-term freshwater fish assemblages and environmental variation in the Ohio River Basin ORSANCO rotenone collections included 89 species total (2,389,722 individuals). Boat electrofishing collections included 90 species total (651,956 individuals). The complete combined dataset included 135 species (not including hybrids) across 19 families. Annual species richness of the combined dataset, excluding rare taxa (< 5 occurrences total), varied from 31 to 90 and increased with year (r = 0.69, *p* < 0.001). Similar patterns were present for correlations of rotenone or electrofishing collections with year. The species with the 10 highest rank counts in the combined dataset were Gizzard Shad (*Dorosoma cepedianum*), Emerald Shiner (*Notropis atherinoides*), Freshwater Drum (*Aplodinotus grunniens*), Channel Shiner (*Notropis wickliffi*), Channel Catfish (*Ictalurus punctatus*), Threadfin Shad (*Dorosoma petenense*), Skipjack Herring (*Alosa chrysochloris*), Common Carp, Bluegill (*Lepomis macrochirus*), and White Bass (*Morone chrysops*). The trophic trait categories with the highest percent counts were herbivore-detritivores (0.34), invertivores (0.19), invertivore-piscivores (0.33), omnivores (0.36), and piscivores (0.09). Life history classifications resulted in 79 opportunistic, 11 equilibrium, 4 periodic, and 41 intermediate species (Appendix 1).

Land use changes during the 1957-2014 period resulted in increased forest land use with year (r = 0.98, *p* < 0.001) and decreased agriculture land use. The IHA for Louisville resulted in significant temporal trends in hydrology for 10 variables that were in four of the five statistics groups defined in IHA software (Table 1). All hydrology variables except fall rate increased during the period. The IHA for Sewickley resulted in significant hydrologic alterations for 10 variables that were in four of the five statistics IHA groups (Table 1). All hydrology variables except 7-day maximum increased. A PCA of Sewickley IHA variables resulted in four PC axes with the first axis correlated with 1-day to 90-day maximum discharge, the second axis correlated with 7-day maximum and reversals, the third axis correlated with December discharge and reversals, and the fourth axis was correlated with December discharge and reversals (Table 2). A PCA of Louisville IHA variables resulted in four PC axes with the first axis correlated with day to 30-day discharge, rise rate, and fall rate, the second axis correlated with November and December discharge, base flow, and high pulse number, the third axis correlated with September, November, and October discharge and base flow, and the fourth axis correlated with September and December discharge (Table 2).

**TABLE 1.** Hydrologic variables that that exhibited evidence of temporal trends during 1957-2014, for Louisville (top) and Sewickley (bottom) USGS gage data.

| IHA statistics group | Hydrologic variables | R<sup>2</sup> |
| --- | --- | --- |
| **Louisville** |  |  |
| Group 1: Magnitude of monthly water conditions | October discharge | 0.10 |
|  | December discharge | 0.11 |
| Group 2: Magnitude and duration of annual extreme water conditions | Annual minimum 1-day | 0.08 |
|  | Annual minimum 3-day | 0.23 |
|  | Annual minimum 7-day | 0.28 |
|  | Annual minimum 30-day | 0.15 |
|  | Annual minimum 90-day | 0.09 |
| Group 3: Timing of annual extreme water conditions |  |  |
| Group 4: Frequency and duration of high and low pulses | High pulse count | 0.13 |
| Group 5: Rate and frequency of water condition changes | Fall rate | 0.25 |
|  | Rise rate | 0.24 |
| **Sewickley** |  |  |
| Group 1: Magnitude of monthly water conditions | December discharge | 0.08 |
| Group 2: Magnitude and duration of annual extreme water conditions | Annual minimum 1-day | 0.20 |
|  | Annual minimum 3-day | 0.31 |
|  | Annual minimum 7-day | 0.10 |
|  | Annual minimum 30-day | 0.24 |
|  | Annual minimum 90-day | 0.08 |
|  | Annual maximum 7-day | 0.29 |
| Group 3: Timing of annual extreme water conditions |  |  |
| Group 4: Frequency and duration of high and low pulses | Low pulse count | 0.13 |
|  | Low pulse duration | 0.23 |
| Group 5: Rate and frequency of water condition changes | Number of reversals | 0.13 |

**Table 2.** Principal components analysis loadings greater than 0.3 for four axes from IHA using data from the Sewickley and Louisville USGS gaging stations

233 **Sewickley, PA USGS gaging station**
| Variable | PCs1 | PCs2 | PCs3 | PCs4 |
| --- | --- | --- | --- | --- |
| 1-day minimum | 0.41 |  |  |  |
| 3-day minimum | 0.42 |  |  |  |
| 30-day minimum | 0.40 |  |  |  |
| 90-day minimum | 0.32 |  |  |  |
| 7-day maximum |  | -0.67 |  |  |
| December |  |  | 0.63 | -0.73 |
| Low pulse number |  |  |  |  |
| Reversals |  | 0.59 | 0.59 | 0.45 |
| Variation explained | 54% | 17% | 8% | 4% |

**Louisville, KY USGS gaging station**
| Variable | PCL1 | PCL2 | PCL3 | PCL4 |
| --- | --- | --- | --- | --- |
| 1-day minimum | 0.31 |  |  |  |
| 3-day minimum | 0.34 |  |  |  |
| 7-day minimum | 0.34 |  |  |  |
| 30-day minimum | 0.34 |  |  |  |
| September |  |  | 0.36 | 0.41 |
| October |  |  | 0.56 |  |
| November |  | 0.36 | -0.40 |  |
| December |  | 0.37 |  | 0.68 |
| Base flow |  | -0.50 | -0.36 |  |
| Hi pulse number |  | 0.31 |  |  |
| Rise rate | 0.31 |  |  |  |
| Fall rate | -0.31 |  |  |  |
| Variation explained | 54% | 16% | 10% | 5% |

### Fish assemblage composition and environmental correlates through time: 1957-2005 rotenone surveys

No single trophic category or life history strategy count was correlated with year, however ordinations did provide evidence of temporal trends. The NMDS analysis using taxonomic counts resulted in a final stress of 0.09 and two axes that explained 83 % of variation (Fig 2). The second NMDS axis was significantly correlated with year (r = 0.64, *p* < 0.001, Fig 2), resulting in a directional trajectory. The NMDS analysis of trophic counts resulted in a final stress of 0.07 and two axes that explained 88 % of variation (Fig 2). The second NMDS axis was significantly correlated with year (r = − 0.41, *p* = 0.02, Fig 2), resulting in a directional trajectory. The NMDS analysis using life history strategies resulted in a final stress of 0.07 and two axes that explained 87 % of variation (Fig 2). Neither life history strategy NMDS axis was significantly correlated with year. The RDA analyses of taxonomic, trophic and life history composition of assemblages did not result in significant ordinations with hydrology and landuse variables.

**FIG 2.**
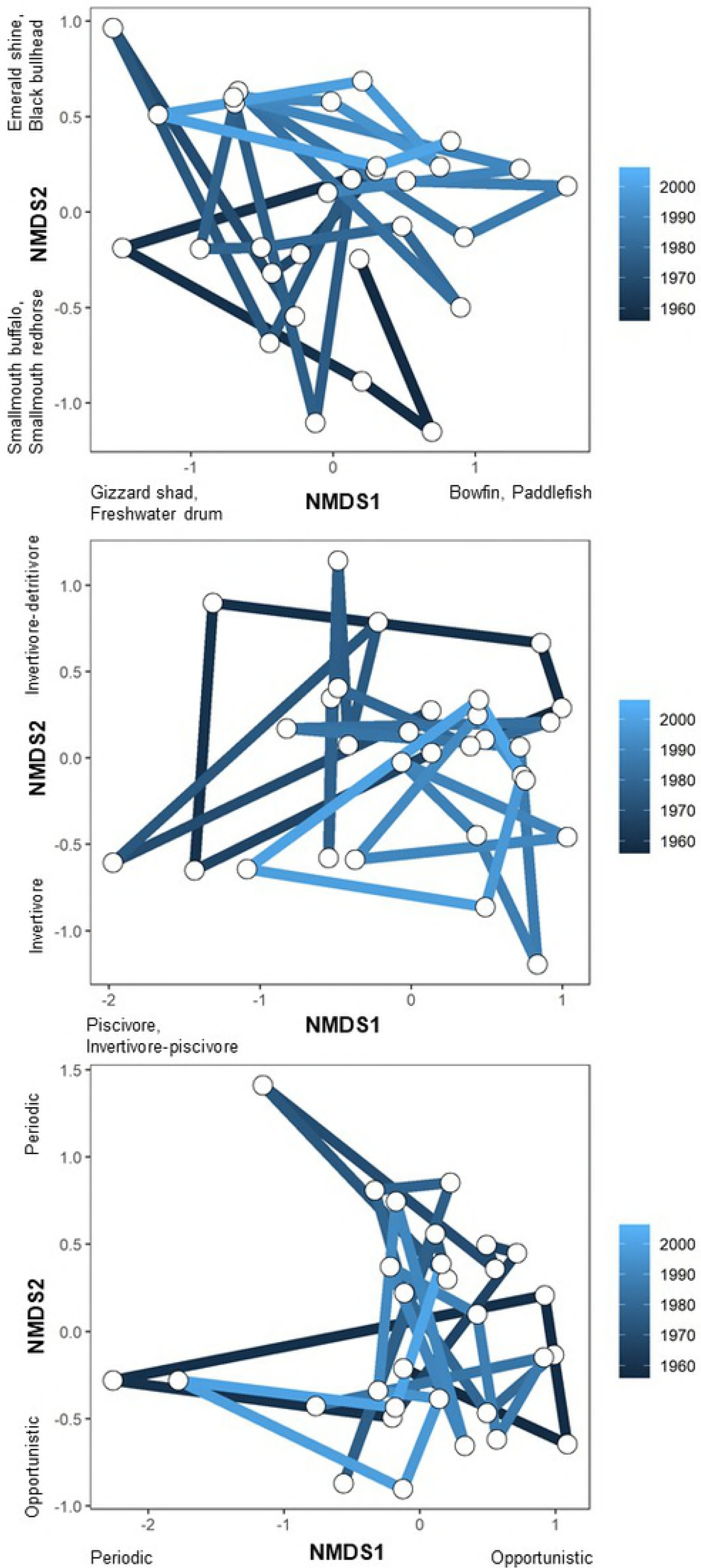
Nonmetric multidimensional scaling ordination of rotenone collections: taxonomic counts (top), trophic counts (middle), and life history strategist counts (bottom). Final stress values were 0.09, 0.09, and 0.07 from top to bottom. Lines connecting years represent temporal trajectories. Highest loading variables are listed on axes.

### Fish assemblage composition and environmental correlates through time: 1990-2014 electrofishing surveys

Trophic categories that increased in count through time included herbivore-detritivores (r = 0.58, *p* = 0.003), invertivores (r = 0.45, *p* = 0.03), invertivore-piscivores (r = 0.51, *p* = 0.01), and omnivores (r = 0.47, *p* = 0.02) (Fig 3). No other trophic trait counts or life history strategies were significantly related to year. NMDS analyses revealed temporal directional trajectories for taxonomic and trophic composition, but not life history composition of assemblages. NMDS analysis of taxonomic composition of assemblages through time resulted in two axes that explained 89% variation (final stress = 0.06; Fig 4). The first NMDS axis was significantly correlated with year (r = 0.89, *p* < 0.001; Fig 4). NMDS analysis of trophic composition of assemblages through time resulted in two axes that explained 86% of variation (final stress = 0.05; Fig 4). The first NMDS axis was significantly correlated with year (r = − 0.89, *p* < 0.001; Fig 4). Finally, NMDS analysis of life history composition of assemblages through time explained 98 % of variation (final stress = 0.01; Fig 4). Neither NMDS axis was significantly correlated with year.

**Fig 3.**
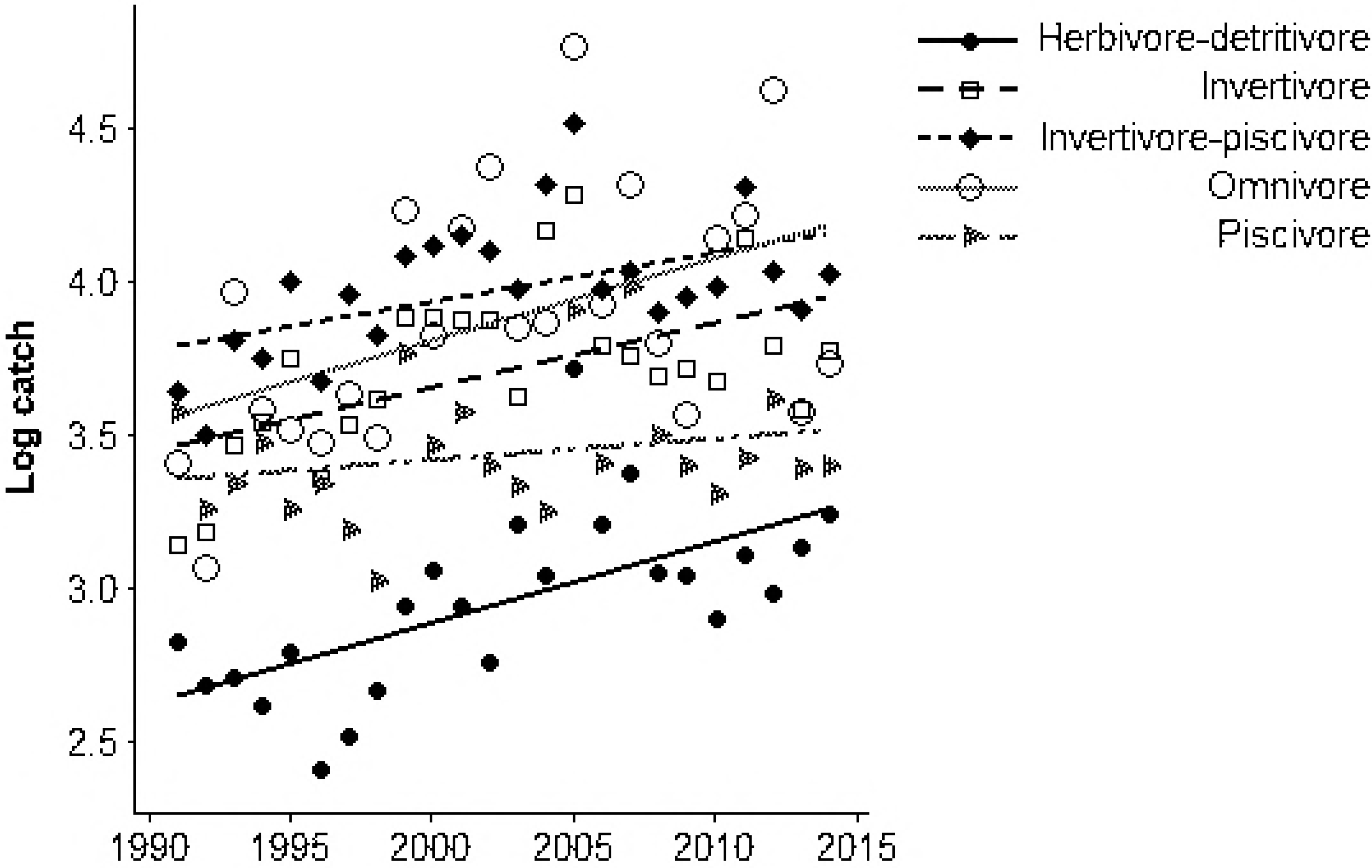
Log-transformed trophic counts by year, for electrofishing collections.

**FIG 4.**
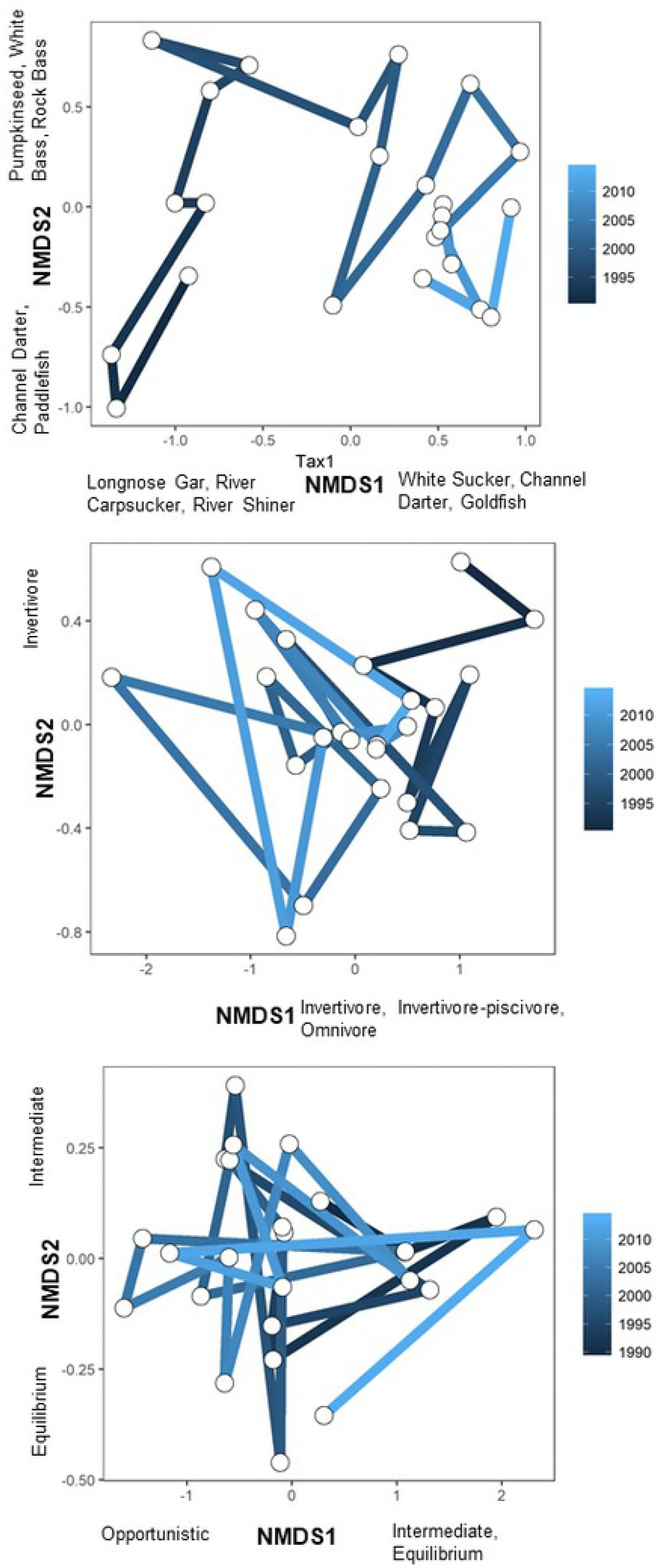
Nonmetric multidimensional scaling ordination of electrofishing collections: taxonomic counts (top), trophic counts (middle), and life history strategist counts (bottom). Final stress values were 0.06, 0.05, and 0.01 from top to bottom. Lines connecting years represent temporal trajectories. Highest loading variables are listed on axes.

RDA of taxonomic composition of assemblages resulted in two significant axes that explained 29% of variation (all canonical axes were significant, *p* = 0.004). Forest land use was significantly correlated with the first RDA axis, and fishes with higher counts in later years were River Carpsucker (*Carpiodes carpio*), Silver Carp, and Spotfin Shiner (*Cyprinella spiloptera*) (Fig 5). Fishes with higher counts in earlier years with decreased forest land use were Goldfish, Channel Darter (*Percina copelandi*), and Slenderhead Darter (*Percina phoxocephala*) (Fig 5). The second RDA axis was significantly correlated with two hydrology PC axes: the first PC axis from the Louisville IHA represented increased minimum discharge, decreased fall rate and increased rise rate (Table 2). Fishes with higher counts in these years were Bighead Carp, Mississippi Silvery Minnow (*Hybognathus nuchalis*), and Spotted Gar (*Lepisosteus oculatus*) (Fig 5). Fishes in lower counts during these years were Smallmouth Bass, Spotted Bass, and Bluegill (Fig 5).

**FIG 5.**
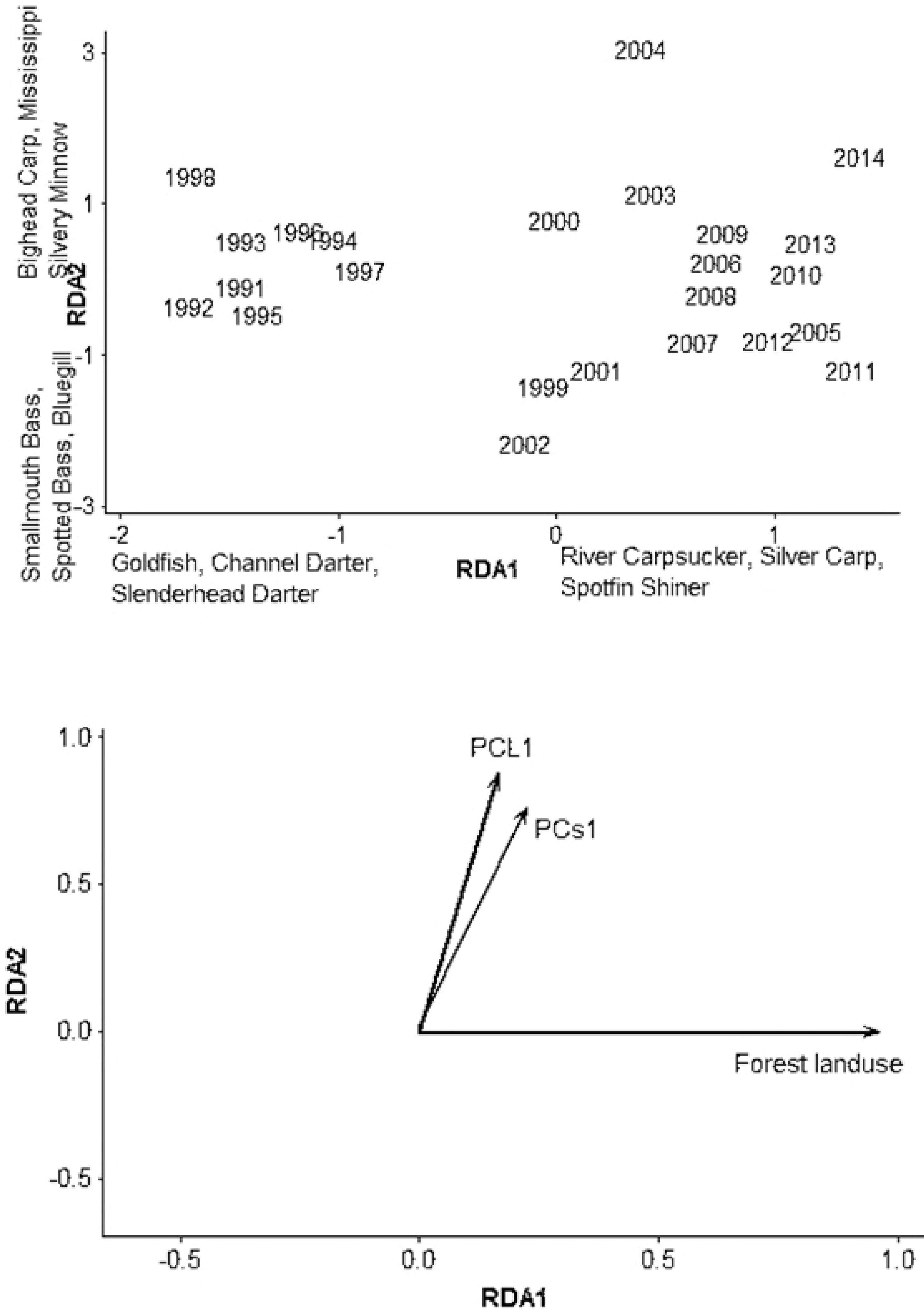
Redundancy analysis using taxonomic counts from electrofishing collections, with significant environmental vectors on bottom. Axes explained 22.4% and 6.4% of variation. PC axes represent altered hydrology variables (loadings are in Table 2). Species or trophic traits with highest loadings on axes are listed.

RDA of trophic composition of fish assemblages resulted in two significant axes that explained 38.8% of variation (all canonical axes were significant, *p* = 0.03). Forest land use was significantly and negatively correlated with the first RDA axis in recent years (Fig 6). Trophic categories with higher counts in these years were herbivore-detritivores. Piscivores and herbivore-detritivores had lower counts during these years. The second RDA axis was strongly correlated with hydrology PC axes. The first PC axis, positively correlated with the second RDA axis, captured increased minimum discharge from both gages (Fig 6, Table 2). The second PC axis, negatively correlated with the second RDA axis, captured years with increased fall discharge, decreased base flows, and increased number of high pulse events from the Louisville gage (Fig 6, Table 2). The second RDA axis was also positively correlated with increased counts of herbivore-detritivores and negatively correlated with counts of piscivores (Fig 6). The RDA of life history composition did not result in a significant ordination.

**Fig 6.**
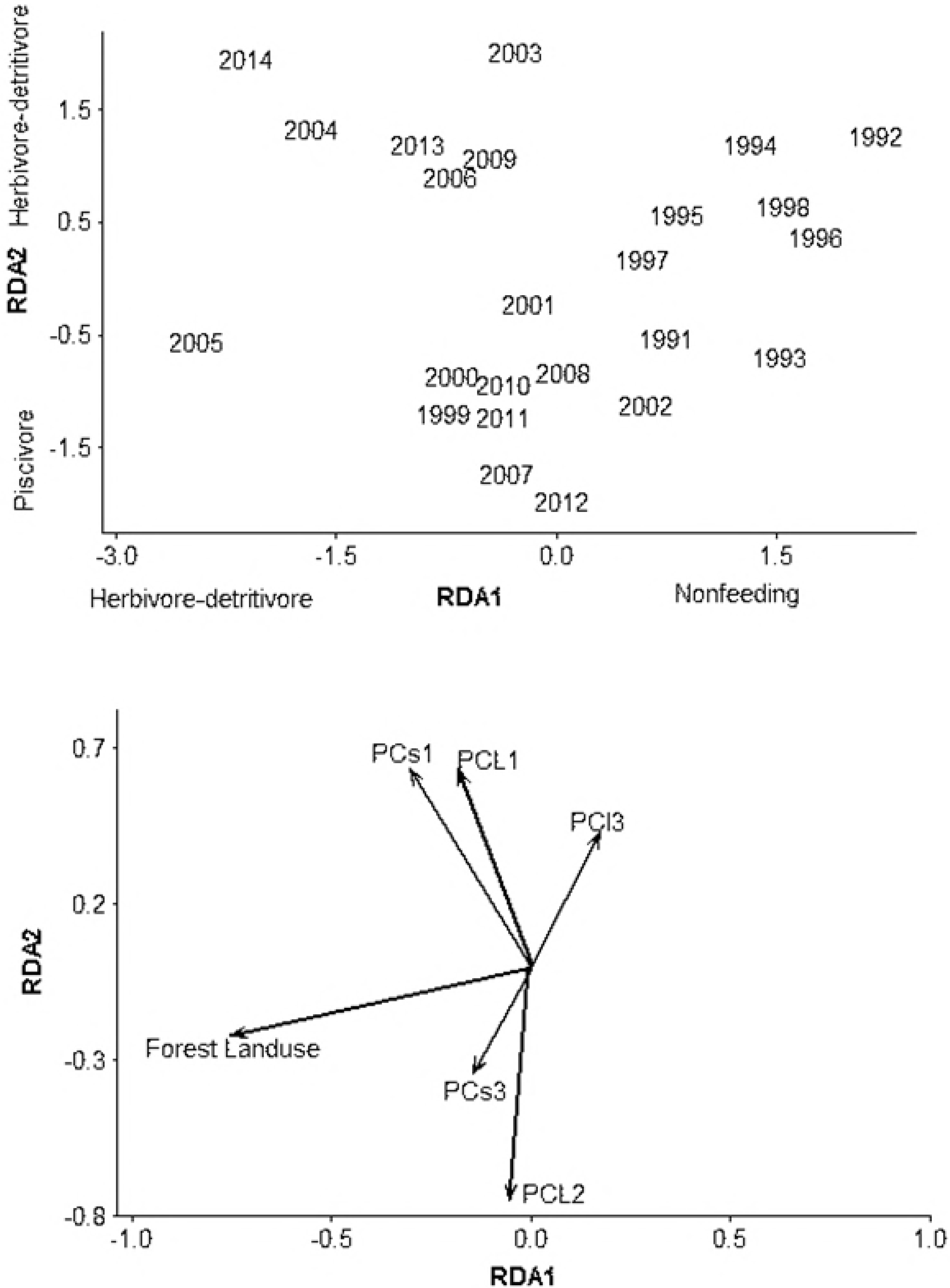
Redundancy analysis using trophic counts from electrofishing collections, with significant environmental vectors on bottom. Axes explained 25.6% and 13.2% of variation. PC axes represent altered hydrology variables. Species or trophic traits with highest loadings are listed on axes.

## Discussion

At the scale of the complete Ohio River mainstem, we found directional shifts in taxonomic and trophic composition of freshwater fish assemblages over multiple decades. Our findings may stem from basin-wide changes over the temporal duration of our study, including water quality improvements following the Clean Water Act, which can influence freshwater assemblages, food webs, and the environmental selection pressures that influence them. Additionally, the inability to detect a change in life history composition of freshwater assemblages at a basin scale may reflect the importance of local scale in considering hydrologic alteration and its effects on freshwater fish assemblages. Our results differed for rotenone and electrofishing data, likely because the rotenone collections were at fewer sites and were not made in all years. In addition, biases in each of these collection methods likely differ among taxa and habitats based upon responses to boat electrofishing or presence in lockchambers.

Temporal trends of taxonomic composition of fish assemblages in the Ohio River Basin We found increases in species richness and counts over the duration of this study throughout the Ohio River Basin. Thomas et al. [11] utilized an index of well-being (MIWB) on this same dataset and reported an increase in count of sensitive taxa and biomass score for fish assemblage quality. Our results and Thomas et al. [11] provide evidence that management of watersheds for improved water quality may influence fish assemblages and can be detected at a basin scale. Similar patterns were found for the Illinois River fish assemblage responses to land use and hydrologic alteration, where recovery appeared to result after the Clean Water Act [40].

The commercial fish catch yield from the Illinois River declined drastically following construction of navigation dams and locks in the 1930s, compared to historic catches, likely from combined effects of agriculture and hydrologic alteration [40]. Commercial fishing has also likely affected the composition of freshwater fish assemblages in the Ohio River Basin over the course of the study. Paddlefish (*Polyodon spathula*), Silver Carp, Bighead Carp, and three catfish species are current Ohio River commercial fisheries [41,42], and commercial fishing likely had an effect on fish assemblages in the past 50 years. Commercial fishing records are sparse due to lack of reporting, and detailed analysis of commercial fishing effects are difficult. Paddlefish accounted for 85% of the 2008 commercial harvest of Ohio River fishes in Indiana and were the most valuable fishery due to egg harvest [43]. In the Indiana section of the Ohio River in 2000, approximately 16,000 kg of catfish were harvested commercially, but by 2014 harvest decreased to approximately 4,500 kg. Size limits for these catfish allowed sexually immature fish to be collected after 2000. Reduced counts of omnivores (including catfish), detected in this study, likely contributed to the reductions in commercial catch rates for catfish.

### Environmental correlates of changes to trophic composition of fish assemblages

We found increased piscivore count (electrofishing collections) associated with increased lentic conditions that occurred after lock and dam construction [20], and we hypothesize that increased phytoplankton productivity may be the mechanism (although planktivore count did not increase). Enhanced phytoplankton productivity can result in bottom-up effects, with increased count of consumers [44] and piscivores [45]. We assume that decreased turbidity occurred with decreasing agricultural land use in the Ohio River Basin over the study period may also have indirect effects on piscivores that capture prey using vision. This is in contrast to invertivores such as catfish that rely on tactile or taste perception and thrive in turbid habitats [46].

Bowes [23] found a 10-year increase in carbon from terrestrial plants (C3) in tissues of invertivore and piscivore fishes in 1960, that coincided with increased forest land use. Phytoplankton contributions to fish tissue covaried less predictably with terrestrial C3 during these time periods [23]. We suggest increased forest land use in the watershed during this period resulted in increased secondary food sources, partial restoration of natural hydrology, and decreased turbidity. Sediment storage upstream of dams and invasive *Dreissena* spp. bivalve molluscs may contribute additionally to decreased turbidity [47]. These turbidity modifications could result in an increased ability of prey to avoid visual predators, facilitating niche development and reduced feeding specialization of consumers. Following an increase in turbidity during early settlement of the Ohio River Basin associated with forest cutting and wetland drainage, turbidity has decreased in the second half of the 21^st^ century, likely due to the conversion of agriculture to forested lands [11].

Altered hydrology variables representing minimum mean discharge and rise and fall rates were correlated with fish composition. These axes were also correlated with decreased counts of piscivores and planktivores, and increased counts of herbivore-detritivores. Increased discharge variability can have negative effects on fishes through life cycle disruption, altered assemblages, and loss of sensitive species [48]. We do not find an obvious mechanistic explanation for these correlations.

Studies at smaller scales (e.g., the Wabash River) offer a comparison to trophic composition of freshwater fish assemblages and environmental attributes of the Ohio River Basin reported here. Anthropogenic influences on the Wabash River and its watershed include hydrologic alterations, land use, historical industrial wastes [49], and the Asian carp invasion in the 1990s [50]. Pyron et al. [50] suggested that changes in Wabash River watershed agriculture nutrient management and Asian carp invasion were contributors to observed fish assemblage variation. These changes include decreased planktivore/detritivores and omnivores, and increased benthic invertivores [51]. The Ohio River watershed has undergone similar agricultural land use modifications, but some differences from the Wabash River remain (e.g., the dominant land use changed from rowcrop agriculture to forest). Intensive rowcrop agricultural land use can influence abundances of generalist fishes (i.e., invertivores) that do not require specialized habitats and prey through nutrient and hydrologic modifications. This may explain trends in trophic composition in the Ohio River basin during this study, including omnivore and invertivore-piscivore dominance during the entire time period.

In addition to changes in trophic composition of fish assemblages found in this study, intraspecific changes may also be occurring that are not detectable at the resolution of static, species-specific trait data. Delong & Thoms [20] identified major changes in carbon sources, and increased variation in mean nitrogen stable isotope ratios of fishes, following flow modification of the Ohio River in 1950-1955. Bowes [23] found decreased trophic position of Ohio River fishes over the past century. Piscivores may switch from traditional planktivore prey (Gizzard Shad) to larval fish, zooplankton, or benthic invertebrates following decreased turbidity; this provides a potential explanation for recent decreased trophic level in piscivores. Bowes [23] showed that Ohio River fish species were more closely packed in isotopic niche space after dams were installed. Contributions of algae to fish tissue carbon increased with dam construction, and the contribution of terrestrial carbon sources decreased. The mean trophic position of Ohio River fishes also decreased following dam construction. Bowes [23] interpreted the causes as a reduction in the relative amount of shallow areas where light can reach the benthos, and thus a reduction in benthic algal productivity. This likely caused a shift from benthic to phytoplankton production, decreased secondary productivity of benthic invertebrates, and decreased trophic position of invertivore and piscivore fishes.

Primary flow alterations of the Ohio River are from dams constructed to enable lockchamber use for barges, resulting in modification of the channel from flowing to lentic conditions [11]. Over the period of the study, we detected signals of increased variability in discharge, decreased seasonality, and generally less stable discharge regimes. We thus expected increased opportunistic and intermediate strategists and decreased equilibrium strategists with decreased stability in discharge regimes in the Ohio River. However, we did not detect changes in counts of life-history strategies through time at the scale of the Ohio River Basin. Fish assemblage variation in the Ohio River and other rivers likely have different patterns at different locations, and at different spatial scales [52]. Indeed, changes in hydrology differed to some degree between the two gages. In this case, our study design relating data from two stream gages to assemblage data throughout the basin revealed trends in taxonomic and trophic composition through time, but it may not reveal the scale at which flow regimes filter for life history strategies. Furthermore, changes in trophic composition through time have stronger hypothesized linkages to landuse and other basin-wide changes than hydrology, which may have stronger mechanistic links to taxonomy and life history. Previous studies supporting relationships between life history strategies and flow regimes have linked assemblages with stream flow data at more local scales (e.g., within stream reaches) rather than across an entire basin (e.g., [27]). At large spatial scales, the ability of hydrology alone to explain life history composition in fish assemblages is likely limited [53].

## Conclusions

We found detectable, directional changes through time in taxonomic and trophic composition of freshwater fish assemblages throughout the Ohio River Basin. Furthermore, we found evidence of relationships between environmental attributes (land use, hydrology) and taxonomic and trophic composition of assemblages. However, we did not find detectable temporal changes in life history composition of assemblages at a basin scale. Future land use modifications, climate change, and altered biotic interactions could continue to contribute to complex and directional patterns of change in freshwater fish assemblages in the Ohio River. Continued efforts to incorporate spatial and temporal scales will help reveal patterns of fish assemblage composition and its environmental correlates across scales, from basins to reaches.

## Acknowledgements

We are grateful to ORSANCO for data access and Jeff Thomas for his assistance in this manuscript. We have no conflicts of interest as no funding was used for this research.

## Authors Contributions

Study design, data analysis, writing: MP.

Study design, writing: MCM.

Data analysis, writing: RCS, NC, CCA.

## Supporting information

### Figure legends

**S1 FIG 1** The Ohio River Basin showing locations of navigational dams (bars) and two USGS gaging stations (open circles). Locations for fish sampling are described by Thomas et al. [11].

**S2 FIG 2** Nonmetric multidimensional scaling ordination of rotenone collections: taxonomic counts (top), trophic counts (middle), and life history strategist counts (bottom). Final stress values were 0.09, 0.09, and 0.07 from top to bottom. Lines connecting years represent temporal trajectories. Highest loading variables are listed on axes.

**S3 FIG 3** Log-transformed trophic counts by year, for electrofishing collections.

**S4 FIG 4** Nonmetric multidimensional scaling ordination of electrofishing collections: taxonomic counts (top), trophic counts (middle), and life history strategist counts (bottom). Final stress values were 0.06, 0.05, and 0.01 from top to bottom. Lines connecting years represent temporal trajectories. Highest loading variables are listed on axes.

**S5 FIG 5** Redundancy analysis using taxonomic counts from electrofishing collections, with significant environmental vectors on bottom. Axes explained 22.4% and 6.4% of variation. PC axes represent altered hydrology variables (loadings are in Table 2). Species or trophic traits with highest loadings on axes are listed.

**S6 FIG 6** Redundancy analysis using trophic counts from electrofishing collections, with significant environmental vectors on bottom. Axes explained 25.6% and 13.2% of variation. PC axes represent altered hydrology variables. Species or trophic traits with highest loadings are listed on axes.

**S7 File. APPENDIX 1** Life history classifications based on Mims & Olden [27] and Perkin et al. [4]. Strategy weights are log-transformed and classified into opportunistic (Opp), periodic (Per), and equilibrium (Equ), and hard classification (Class) includes intermediate classifications

